# Effects of metabotropic 2/3-glutamate receptor activation and inactivation on activity-based anorexia model in mice

**DOI:** 10.64898/2026.02.06.704365

**Authors:** Tuomas Varjelus, Atte Oksanen, Laura Kaljala, Maria Ryazantseva, Teemu Aitta-aho

## Abstract

*Anorexia nervosa* is an eating disorder disproportionately found in female human teens and young adults. It is often resistant to treatment, has a significant chance of relapse and is more lethal than other eating disorders, such as *bulimia nervosa* or Avoidant/Restrictive Food Intake Disorder (ARFID). There is no specific medication for the treatment of *anorexia nervosa*. Treatment consists of psychosocial means, psychotherapy, psychoeducation, and nutritional counseling. Medication is usually used for treating comorbidities such as anxiety or to decrease obsessive-compulsive tendencies. These medications cannot help the patient regain weight or treat core symptoms. Metabotropic 2/3-glutamate (mGluR2/3) receptor agonist (LY379268) and antagonist (LY341495) are promising pharmacological agents to treat psychiatric disorders. Both agonists and antagonists have been reported to have anxiolytic effects in different animal models of anxiety, while antagonists have shown antidepressant-like effects in preclinical studies. The activity-based anorexia (ABA) paradigm is used to model *anorexia nervosa*. It consists of giving mice access to a running wheel and restricting their feeding time. This causes mice to exercise more than mice without feeding time restriction and to eat less than mice without access to a moving running wheel. In this study, we subcutaneously injected female ABA model mice with a metabotropic 2/3-glutamate receptor agonist (LY379268) and antagonist (LY341495) in two experiments. Both compounds exacerbated weight loss by decreasing food intake as well as increasing physical activity. It can be concluded that the manipulation of mGluR2/3 receptors is detrimental for the ABA model and likely for *anorexia nervosa* as well.

**Highlights:**

- mGluR2/3 agonist LY379268 decreases food intake and body weight of the ABA model
- mGluR2/3 antagonist LY341495 decreases food intake and body weight of the ABA model
- Both agonist and antagonist produce the effect within 48 hours
- Both the agonist and antagonist are detrimental to the ABA-model

## 1. Introduction

*Anorexia nervosa* (AN) is an eating disorder characterized by fear of gaining weight, reduction in food intake, and hyperactivity that is disproportionately found in biological female human teens and young adults, as supported by population studies. Nevertheless, it is worth mentioning that the condition can also affect the male population, and diagnostic and historical biases can contribute to the gap in detected prevalence (Bulik et al., 2006; Capuano et al., 2025; Micali et al., 2013). AN is often resistant to treatment, has a significant chance of relapse and is more lethal than other eating disorders, such as *bulimia nervosa* or Avoidant/Restrictive Food Intake Disorder (ARFID). There is no specific medication for the treatment of AN, although antidepressants and antipsychotics are used to treat comorbid psychiatric symptoms such as anxiety, depression, or obsessive-compulsive tendencies, but not the disorder itself (Bauschka et al., 2023; Hartman, 1995; Walsh et al., 2006). The main available treatment consists of psychotherapy, psychoeducation, and nutritional counseling.

The activity-based anorexia (ABA)-model in rodents is well established in preclinical research and is the most often used model of AN (Foldi, 2024; Klenotich and Dulawa, 2012). The ABA model mimics well the biological hallmarks of AN, such as prioritizing activity over sustenance when facing starvation. Moreover, it replicates the endophenotype of AN, characterized by alterations in neural circuits and neurochemistry. It can, as is accepted in the field, provide a mechanistic understanding and rapid translation of novel treatment strategies for AN (Foldi, 2024). The model protocol includes the introduction of a running wheel for unrestricted physical activity, with food available for a limited time daily for the ABA group. The control groups include mice that have a running wheel and are not restricted in food intake (“Wheel” group), and mice that are restricted in food intake time for the same period as the ABA group and have a locked wheel with no option to run (“Food restriction” group). As a result, mice exposed to the ABA protocol run more than mice without food restriction and eat less than mice without a moving running wheel (Klenotich and Dulawa, 2012). Importantly, in the ABA model, food is limited only by time, not quantity. Without access to a running wheel, the control animals quickly learn to eat enough food within the limited time window, while ABA-treated mice voluntarily reduce food intake as soon as anorexia symptoms emerge. The model has no negative off-target effects of genetic models and replicates a key aspect of AN in voluntary reduction in food intake (Foldi, 2024).

Metabotropic glutamate receptors (mGluRs) are promising targets for treating brain disorders because they can be modulated to fine-tune synaptic activity and protect neurons. Unlike ionotropic glutamate receptors (NMDA and AMPA), mGluRs modulate rather than mediate synaptic transmission. This is a significant advantage, as mGluR-targeting drugs may have fewer side effects than directly blocking excitatory signals (Alborghetti et al., 2025; Dogra and Conn, 2021a, 2021b; Hovelsø et al., 2012; Maksymetz et al., 2017; Masilamoni and Smith, 2018). There are three main groups of metabotropic glutamate receptors. Receptors mGluR2 and mGluR3 belong to Group II and are coupled to the inhibition of adenylyl cyclase activity through G-proteins of the G_i_/G_0_ type (Dogra and Conn, 2021a; Hovelsø et al., 2012). Group II metabotropic glutamate receptors can be located pre-or postsynaptically, but mGluR2 is predominantly presynaptic, where it mainly regulates transmitter release, while mGluR3 is found both presynaptically and postsynaptically, including in glial cells. Both receptors generally act to reduce glutamate or GABA release and dampen neuronal excitability through intracellular pathways such as inhibition of adenylate cyclase, modulation of mTOR signaling, regulation of ion channels, and induction of long-term depression (Chen et al., 2001; Haroon et al., 2017; Hong et al., 2009; Hovelsø et al., 2012; Roh et al., 2024). mGluR2 and mGluR3 show overlapping but distinct expression patterns in the rodent brain. Both receptors are widely distributed in corticolimbic and subcortical structures, but their relative abundance differs. mGluR2 shows high expression in limbic and associative regions, particularly in the hippocampus (dentate gyrus, CA3, and to a lesser extent CA1), basolateral amygdala, prefrontal cortex, and other frontal cortical areas, thalamus and specific relay nuclei, and olfactory regions, including the olfactory bulb and piriform cortex. mGluR3 has a broader and more diffuse distribution, with prominent expression in the cortex, widespread neocortical layers, hippocampus (CA1 and CA3), striatum, cerebellum (including molecular and granular layers), thalamus, and hypothalamus (McOmish et al., 2016; Ohishi et al., 1994; Testa et al., 1994; Woo et al., 2022; Wright et al., 2013).

Agonism of group II metabotropic glutamate receptors demonstrates potential as an antipsychotic-like and procognitive treatment (Dogra and Conn, 2022; Takamori et al., 2003). The agonists have consistently shown anxiolytic activity in a wide range of animal models of anxiety (Ferraguti, 2018; Linden et al., 2005; Schoepp, 2024). Inhibiting mGluR2/3 receptors offers a novel, potentially safer pathway to achieve rapid antidepressant effects (Chaki, 2020; Onisiforou et al., 2023; Potter et al., 2020). It was previously determined that the mechanism underlying the antidepressant-relevant actions of ketamine converges with mGluR2 signaling (Zanos et al., 2019). Additionally, the benefit of metabotropic 2/3-glutamate receptor agonists over ketamine is that they are not addictive or sedative but offer similar benefits through the glutamate pathways (Chaki, 2020; Sos et al., 2013).

Classic antidepressants, such as SSRIs, have shown inconsistent results in clinical use to treat AN, suggesting that they are ineffective at the acute stage but can be beneficial after body weight restoration (Walsh et al., 2006). Ketamine is also considered a potential treatment for AN (Hermens et al., 2020). It is suggested that ketamine may reduce adult females’ vulnerability to ABA and may protect women from AN relapse by reducing hyperactivity (Dong and Aoki, 2025a). The action of subanaesthetic doses of ketamine in normalizing neurotransmission and alleviating symptoms is associated with the downregulation of NMDA receptors (Mitchell et al., 2023; Schalla and Stengel, 2019). Antipsychotics can also be used to treat some of the symptoms of AN (Bauschka et al., 2023; Han et al., 2022; Hernández Sánchez et al., 2016), with olanzapine showing promising results (Han et al., 2022). As anxiety is often comorbid with AN and anxiety as a trait is associated with AN (Sternheim et al., 2024; Udo and Grilo, 2019), the anxiolytic effect of potential treatments, including antipsychotics, can be beneficial for some patients (Bauschka et al., 2023).

Based on all that was said above, in this study, we conducted experiments to evaluate the effects of metabotropic 2/3-glutamate receptor agonist and antagonist on the anorexia phenotype in female ABA model mice. In particular, we addressed the acute stage of anorexia symptom development, when voluntary food intake was significantly reduced despite high physical activity.

## 2. Materials and methods

### 2.1. Animals

Female C57BL/6JRcc mice aged 6 to 10 weeks were used in both experiments. They were born and kept at a reversed circadian rhythm with lights off from 6 to 18. Before the start of the experiment, the mice received food and water ad libitum. The mice were bred from the University of Helsinki Laboratory Animal Center’s stock (originally from Envigo/<u>Inotiv, Inc</u>.). All animal experiments were performed following the University of Helsinki Animal Welfare Guidelines and approved by the National Animal Experiment Board of Finland (license: ESAVI/2587/2024).

### 2.2. Activity-based anorexia model

The mice were housed in separate individually ventilated cages (IVCs) with low-profile wireless running wheels (ENV-047, Med Associates Inc.). The mice were first habituated to the environment for a week before the introduction of feeding time restriction and injections. Feeding time was restricted to 1.5 hours per day, starting 30 minutes after injection, in the mornings at approximately 9:30 AM. Animals were removed from the experiment when they reached their endpoint weight, defined as a 25% decrease from the original body weight. The mice were sacrificed using CO_2_ and cervical dislocation.

### 2.3. Experiment 1

In this experiment (Fig. 1A), we tested the metabotropic glutamate receptor 2/3 agonist LY379268 in the ABA model. Mice were grouped into 6 groups, each consisting of 4 mice. The first group, “Food restricted + saline”, had their feeding time restricted to 1.5 hours per day and was injected with saline (0.9% NaCl, 10 ml/kg). Their running wheels were locked. The second group, “Food restricted + LY,” had their feeding time restricted to 1.5 hours per day but were injected with LY379268 (3 mg/kg in saline 0.9% NaCl, 10 ml/kg). Their running wheels were also locked. The third group, “Wheel + saline”, had no restrictions on feeding time and was injected with saline (same dose as the first group). Their running wheels moved normally. The fourth group, “Wheel + LY”, underwent the same treatment as the third group, but with LY injections (same dose as the second group). The fifth and sixth groups were exposed to the ABA protocol: feeding time was restricted to 1,5 hours per day, and the wheels were not locked to allow voluntary running. The fifth group, “ABA + saline”, was injected with saline, and the sixth group, “ABA + LY”, was injected with LY. The same dosages were used as for the other groups.

**Fig. 1.**
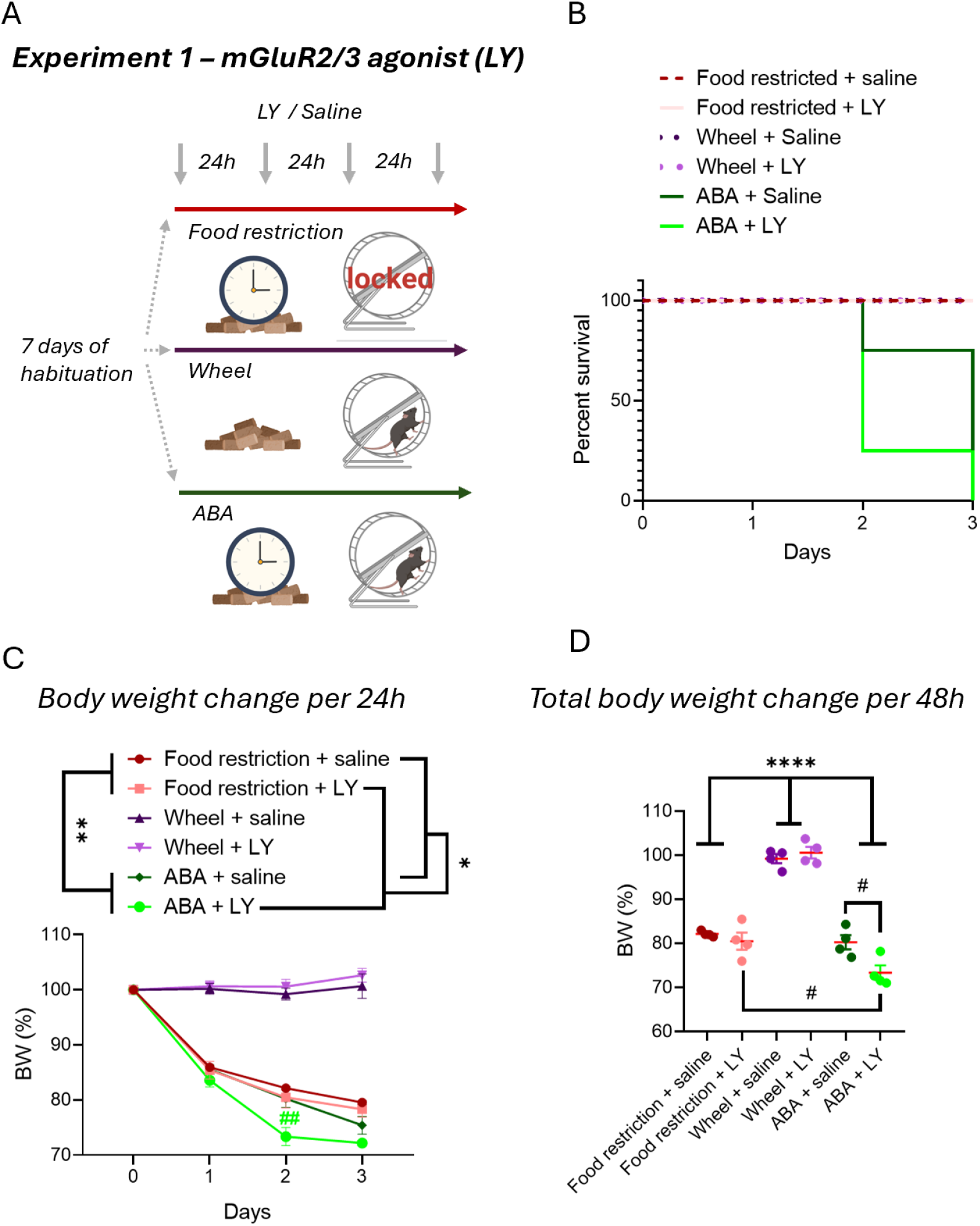
Effects of daily administration of the mGluR2/3 agonist on survival and body weight in the activity-based anorexia (ABA) model and controls. **A**. Scheme of Experiment 1 with mGluR2/3 agonist LY379268 (LY) or saline administration to ABA and control groups (see methods). **B**. Survival plot: mice reaching the endpoint due to body weight loss (4 mice per group). **C**. Body weight (BW) changes every 24 h (days) after the start of the ABA protocol (N=4, mixed effect model (REML), food restriction vs. ABA: F (1, 12) = 11.11, **p=0.006; LY vs saline in food restriction and ABA groups: F (1, 12) = 5.626, *p=0.0353; data shown as the mean ± S.E.M). **D**. Average body weight loss per first 48 hours in groups during the agonist trial. (N=4, two-way ANOVA, ABA and food restriction vs wheel: F (2, 18) = 149.3, ****p<0.0001; post hoc Tukey’s multiple comparisons test, ABA + LY vs ABA + saline: ^#^p=0.0293; ABA + LY vs. food restriction + LY: ^#^p=0.0228, ABA + LY vs. food restriction + saline, ^##^p=0.0041, data shown as the mean ± S.E.M).

The mice were weighed and injected at 9:00 in the morning. The injections were given *subcutem* under the loose skin behind the neck. They were fed for 1.5 hours per day, starting 30 minutes after injection. The food was weighed before and after feeding. The wireless running wheels continuously measured turns through their software, and the number of turns per 24 hours was recorded. The mice that reached their humane endpoint weight or at the end of the 5-day experiment were sacrificed using CO_2_ and confirmed with cervical dislocation.

### 2.4. Experiment 2

In this experiment (Fig. 3A), we tested the metabotropic 2/3-glutamate receptor antagonist LY341495 on the ABA model. The mice were divided into 2 groups of 6 each. One group, “ABA + LY”, was injected with the metabotropic 2/3-glutamate receptor antagonist LY341495 (1 mg/kg in saline 0,9% NaCl, 10 ml/kg), and the other, “ABA + Saline”, was injected with saline (0,9% NaCl, 10 ml/kg).

### 2.5 Drugs

The metabotropic glutamate receptor 2/3 agonist LY379268 or antagonist LY341495 (Tocris Bioscience) was dissolved in equimolar NaOH and adjusted to a pH of 7 with HCl. Drugs were then diluted in saline (0.9% NaCl). The final LY379268 concentration of the solution was 0.3 mg/ml, and the dose in saline was 3 mg/kg. The final LY341495 concentration in saline was 0.1 mg/ml, and the dose was 1 mg/kg. Drugs were weighed using a scale with an accuracy of one-tenth of a microgram.

### 2.6 Statistical testing

GraphPad Prism 10 (GraphPad Software) was used to analyze the raw data and to plot the graphs. An appropriate statistical test was chosen based on the hypothesis and data properties. Analysis was corrected for multiple comparisons. Tests and statistical data are described in the figures and text.

## 3. Results

### 3.1. Experiment 1

In the first experiment, we aimed to evaluate the effects of the mGluR2/3 agonist LY379268 (LY, agonist) on anorexia symptom manifestation in the ABA model in comparison to mice that were restricted in feeding time or had a running wheel only (N=4 mice per group). (Fig. 1A). The LY agonist was administered once a day before feeding, starting from day 0 of the ABA protocol. The protocol significantly affected mouse survival after the start (75% survival at 48 hours, 25% at 72 hours) compared to “Food restriction” (100% at 72 hours) and “Wheel” (100% at 72 hours) controls (log-rank Mantel‒Cox test, χ^2^=26.25, df=5, p<0.0001) (Fig. 1B). At the same time, LY agonist administration reduced the survival possibility of ABA-treated mice from 75% to 25%, with the majority reaching the body weight endpoint within 48 hours (Fig. 2B and C). Although, in general, body weight loss was more severe in the ABA-treated mice than in the “Food restricted” groups (mixed-effects model (REML), ABA *vs*. food restriction treatment effect: F(1, 12)=11.11, p=0.0060) (Fig. 1C), LY administration worsened the effect (mixed-effects model (REML), LY *vs*. saline treatment: F(1, 12)=5.626, p=0.0353). LY administration did not affect the body weight of the “Wheel + LY” group compared to the “Wheel + Saline” group on any day of assessment (two-way ANOVA, F (1, 6) = 0.5436, p=0.4888) (Fig. 1C). The “ABA” and “Food restriction” groups lost significantly more body weight than the control “Wheel” groups (two-way ANOVA, F (2, 18) = 149.3, p<0.0001). Both saline-treated “Food restricted + Saline” and “ABA + Saline” animals lost approximately 20% of their initial body weight in two days compared to “Wheel + Saline” controls (“Food restricted + Saline”: 82.13 ± 0.323%; “ABA + Saline”: 80.25 ± 1.614%; “Wheel + Saline”: 99.22 ± 1.043%) (Fig. 1D). The LY agonist treatment, however, worsened the loss only in “ABA” animals within this time period, making it worse compared to “ABA + Saline” and to “Food restriction + LY” (“ABA + Saline”: 80.25 ± 1.614%; Food restricted + LY”: 80.50 ± 1.958 %; “ABA + LY”: 73.35 ± 1.651%; “Wheel + LY”: 100.58 ± 1.300%; post hoc Tukey’s multiple comparisons test: “ABA + Saline” *vs*. “ABA + LY”: p=0.0293; “Food restriction + LY” *vs*. “ABA + LY”: p=0.0228; “Food restriction + Saline” *vs*. “ABA + LY”: p=0.0041) (Fig. 1D). There was no significant difference after 48 hours from the start of LY administration in comparison to saline treatment for body weight in the “Food restriction” groups (post hoc Tukey’s multiple comparisons test: “Food restriction + LY” *vs*. “Food restriction + Saline”: p=0.9608). Overall, mGluR2/3 agonism with LY379268 had a detrimental effect on animal survival and body weight loss in the ABA model, with the effect being pronounced within 48 hours after the start of the treatment.

**Fig. 2.**
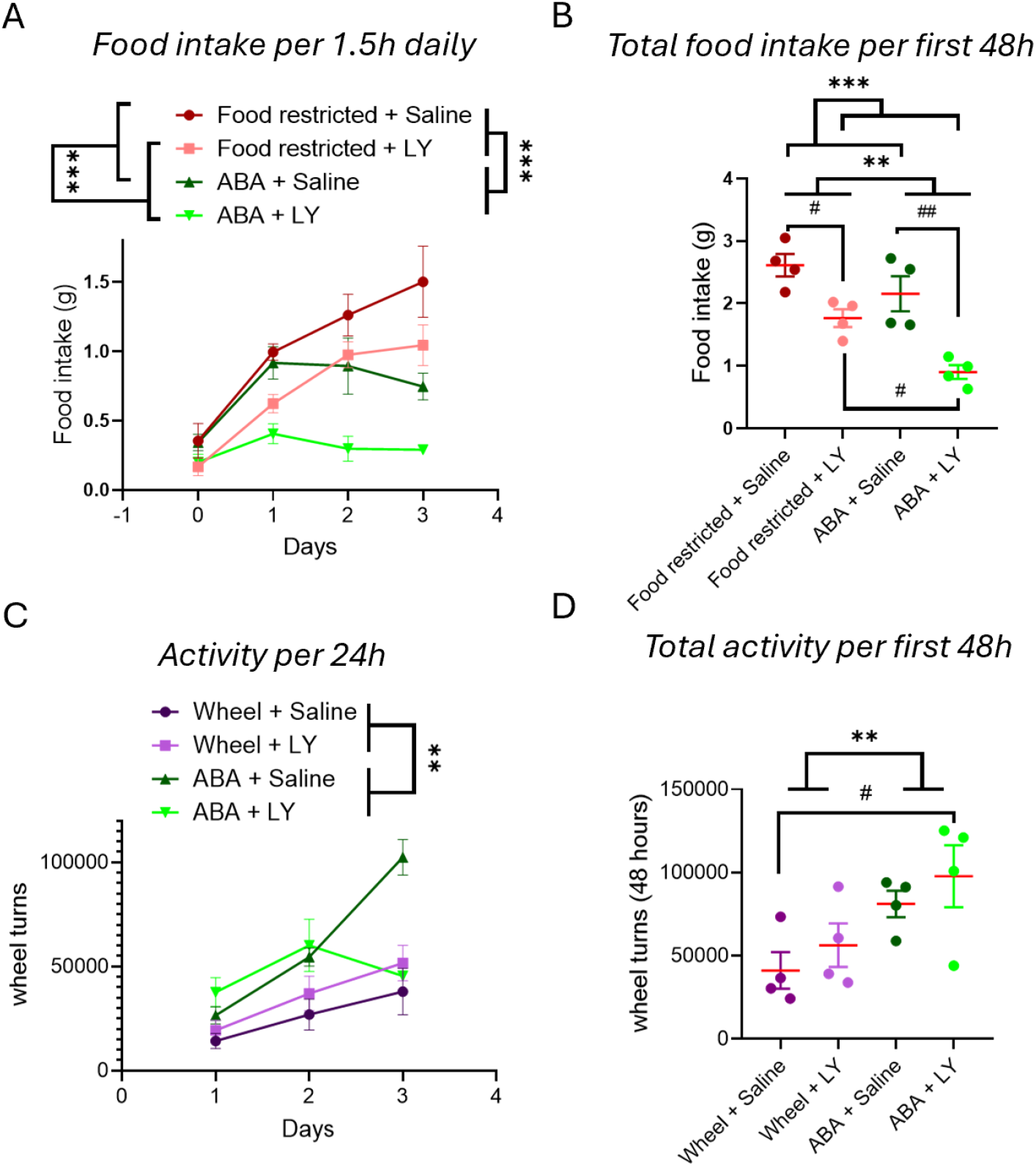
Effects of mGluR2/3 agonism on feeding and activity behaviors in the ABA model and controls. **A**. Food intake per 1.5 hours at 24-hour intervals (days) for the “ABA” and “Food restricted” groups injected with saline or mGluR2/3 agonist (LY) (4 animals per group, “ABA” vs. “Food restriction” treatment: REML, F (1, 12)=21.62, ***p=0.0009, “LY” vs. “Saline” treatment: Mixed effect model (REML), F (1, 12)=23.98, ***p=0.0004; data shown as the mean ± S.E.M.). **B**. Total food intake per 48 hours in “Food restricted” or “ABA” groups injected with saline or LY (N=4, two-way ANOVA, ABA treatment effect: F (1, 12) = 12.19, **p=0.0045, LY treatment effect: F (1, 12) =30.85, ***p=0.0001, post hoc Tukey’s multiple comparisons test, “ABA + LY” vs. “ABA + Saline” ^##^p=0.0032, “Food restriction + LY” vs. “Food restriction + Saline”, p=0.0474, “ABA + LY” vs. “Food restriction + LY”, p=0.0428; data shown as Mean ± S.E.M). **C**. Running activity of ABA and Wheel groups per 24 hours, injected with LY or saline and assessed as wheel turns (N=4, mixed effect model (REML), ABA vs. Wheel treatment effect: F (1, 12) = 13.37, p=0.0033; LY vs. saline treatment effect: F (1, 12) = 0.1076, p=0.7486). **D**. Total activity per 48 hours in wheel turns for mice in the ABA or Wheel groups injected with LY or saline (N=4, two-way ANOVA, ABA treatment effect: F (1, 12) = 9.404, **p=0.0098, LY treatment effect F (1, 12) = 1.439, p=0.2535; post hoc Tukey’s multiple comparisons test, “Wheel + Saline” vs. “ABA + LY”, ^#^p=0.0460; data shown as the mean ± S.E.M.).

**Fig. 3.**
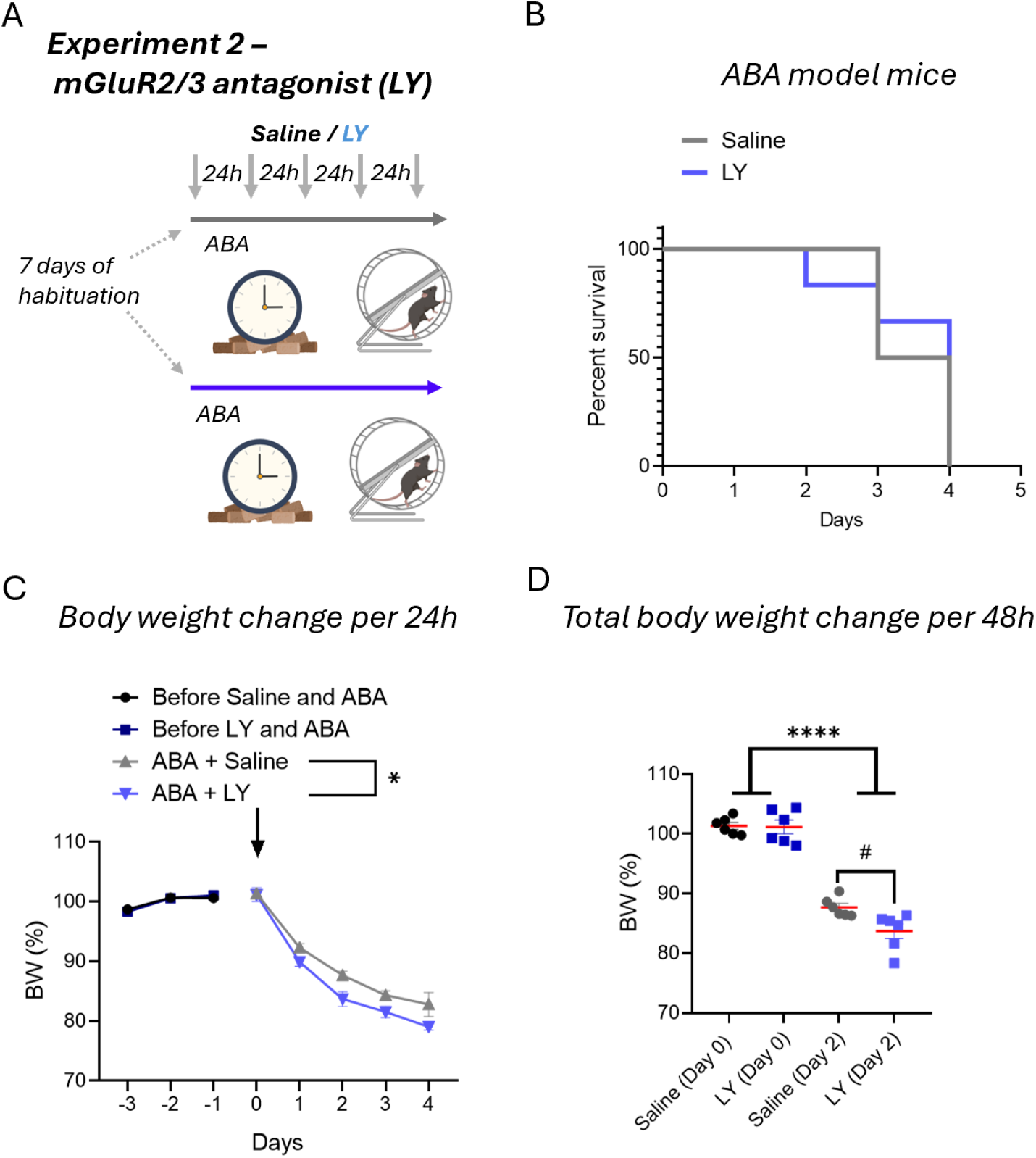
Effects of daily administration of the mGluR2/3 antagonist on survival and body weight in the ABA model. **A**. Scheme of Experiment 2 with mGluR2/3 antagonist LY341495 (LY) or saline administration to ABA groups (see methods). **B**. Survival plot: mice reaching the endpoint due to body weight loss (6 mice per group). **C**. Body weight change per 24 hours for two groups of mice: before and after exposure to the ABA protocol and daily injection with saline or LY. The arrow shows the start of the ABA protocol and pharmacological treatments (N=6, mixed effect model (REML), before the experiment: F (1, 10) = 3.079e-011, p>0.999; LY effect: F (1, 10) = 6.728, *p=0.0268; data shown as the mean ± S.E.M). **D**. Total body weight change of the ABA groups at 48 hours after injection with saline or LY compared to the start of drug administration (day 0) (N=6, two-way RM ANOVA, body weight change: F (1, 10) = 721.5, p<0.0001; post hoc Holm‒Sidak test, saline (day 2) vs. LY (day 2), ^#^p=0.0156, data shown as the mean ± S.E.M.).

The ABA-treated mice consumed significantly less food per 1.5 hours daily than the controls, which was restricted in their feeding time in the same manner (“ABA” *vs*. “Food restriction” treatment: Mixed effect model (REML), F (1, 12) = 21.62, p=0.0006). Daily LY agonist administration reduced food intake in both the “Food restriction + LY” and “ABA + LY” groups compared to saline-injected controls (“LY” *vs*. “Saline” treatment: Mixed effect model (REML), F (1, 12) = 23.98, p=0.0004) (Fig. 2A). Within 48 hours, the ABA-protocol-treated mice ate less than “Food restricted” mice (two-way ANOVA, ABA treatment effect: F (1, 12) = 12.19, p=0.0045) (Fig. 2B). Agonist administration also produced significant effects in both groups (two-way ANOVA, LY treatment effect: F (1, 12) = 30.85, p=0.0001). Within this period, the food intake of the “ABA + LY” group (0.90 ± 0.111 g total) was lower than that of the “ABA + Saline” group (2.16 ± 0.279 g total, post hoc Tukey’s multiple comparisons test, p=0.0032) and lower than that of the “Food restriction + LY” group (1.77 ± 0.142 g total, post hoc Tukey’s multiple comparisons test, p=0.0428). The food intake of the “Food restriction + LY” group, at the same time, was lower than that of the “Food restriction + Saline” group (2.61 ± 0.180 g total, post hoc Tukey’s multiple comparisons test, p=0.0474). Therefore, mGluR2/3 agonism reduced food intake within 48 hours when feeding was time-restricted and accelerated anorexia-like symptoms in the ABA model.

Despite low food consumption, the amount of physical activity in the running wheel was elevated in the “ABA” groups compared to mice with ad libitum access to food (mixed effect model (REML), ABA *vs*. wheel treatment effect: F (1, 12) = 13.37, p=0.0033) (Fig. 2C). The LY agonist did not produce a general significant elevation in activity within the groups when the activity was measured daily and tested for the whole experiment duration (mixed effect model (REML), LY *vs*. saline treatment effect: F (1, 12) = 0.1076, p=0.7486) (Fig. 2C). The accumulated amount of activity (total wheel turns) for 48 hours was higher in the “ABA” groups (Saline: 81141 ± 7967 turns, LY: 97814 ± 18667 turns) than in the “Wheel” groups (Saline: 41192 ± 11029 turns, LY: 56360 ± 13078 turns) (two-way ANOVA, ABA treatment effect: F (1, 12) = 9.404, p=0.0098). Although there was no significant effect of LY on either treated group (two-way ANOVA, LY treatment effect F (1, 12) = 1.439, p=0.2535), the “ABA + LY” group’s activity was elevated compared to that of the “Wheel + Saline” control group (post hoc Tukey’s multiple comparisons test, p=0.0460) (Fig. 2D). Although there was no clear effect of the drug on physical activity, it is not possible to completely exclude that mGluR2/3 agonism contributes to the elevated physical activity in “ABA + LY” mice, as the size of the effect can be low due to malnutrition, exhaustion and life-threatening weight loss.

Taken together, mGluR2/3 agonism produces a detrimental effect on anorexia-like symptoms in the ABA model by strongly contributing to an accumulating reduction in food intake that leads to life-threatening body weight loss.

### 3.2. Experiment 2

In Experiment 2, we aimed to determine whether mGluR2/3 antagonism with LY341495 (LY, antagonist) has a positive effect on the ABA model (Fig. 3A). In this experiment, two groups of mice (N=6 mice per group) were subjected to the ABA protocol and injected with saline or LY antagonist every 24 hours. There was no significant difference in survival rate between ABA-treated mice injected with saline or LY (log-rank Mantel‒Cox test, χ^2^=10.72, df=1, p=0.7433). As in Experiment 1, mice reached the endpoint starting at 48-72 hours into the ABA protocol and pharmacological treatment (Fig. 3B). However, body weight loss progression was significantly higher in LY antagonist-injected mice than in saline-injected mice (mixed effect model (REML), LY effect F (1, 10) = 6.728, p=0.0268). Before the experiment, these two groups showed no difference in body weight changes for 3 days (mixed effect model (REML), group effect: F (1, 10) = 3.079e-011, p>0.999) (Fig. 3C). Within 48 hours of the experiment, the mice lost from 10% to 20% of their body weight (body weight ABA + saline: 87.69 ± 0.651% and ABA + LY: 83.70 ± 1.260% of initial body weight; two-way ANOVA, initial *vs*. after 48 h body weight F (1, 10) = 721.5, p<0.0001) (Fig. 1D). The loss was more severe in LY antagonist-treated animals than in saline-treated animals (post hoc Holm‒Sidak test, p=0.0156) (Fig. 1D). Overall, although mGluR2/3 antagonism did not improve or worsen the survival rate of ABA-treated mice, it significantly contributed to weight loss within 48 hours of daily drug administration.

In line with the body weight loss effect, daily LY antagonist administration significantly reduced food intake in ABA-treated mice (mixed effect model (REML), ABA + LY *vs*. + saline treatment: F (1, 10) = 8.056, *p=0.0176). Before the start of LY antagonist administration and the ABA protocol, these two groups consumed the same amount of food daily (REML, group effect before treatment: F (1, 10) = 0.4005, p=0.5410) (Fig. 4A). Within 48 hours of LY antagonist administration, drug-treated mice consumed significantly less food than saline-treated mice (Saline: 3.87 ± 0.223 g; LY: 2.78 ± 0.320 g; t test, t=2.778, df=10, p=0.0195) (Fig. 4B). Both ABA groups were active in the running wheel, with the LY antagonist-administered group showing more wheel turns during the experiment (two-way ANOVA, LY effect: F (1, 10) = 6.708, p=0.027) (Fig. 4C). Before the start of the ABA protocol and injections, the groups demonstrated no difference in daily activity (two-way ANOVA, group effect: F (1, 10) = 0.2798, p=0.6084). Within 48 hours of LY antagonist administration, mice ran 41394 ± 6763 wheel turns, whereas mice injected with saline ran 27990 ± 3730 turns. Despite the clear trend in the elevation of running activity at the 48-hour distance, the difference was not statistically significant (t test, t=1.736, df=10, p=0.1133) (Fig. 4D). Therefore, 2 days of mGluR2/3 antagonist treatment reduced the food intake of ABA mice, worsening the anorexia phenotype.

**Fig. 4.**
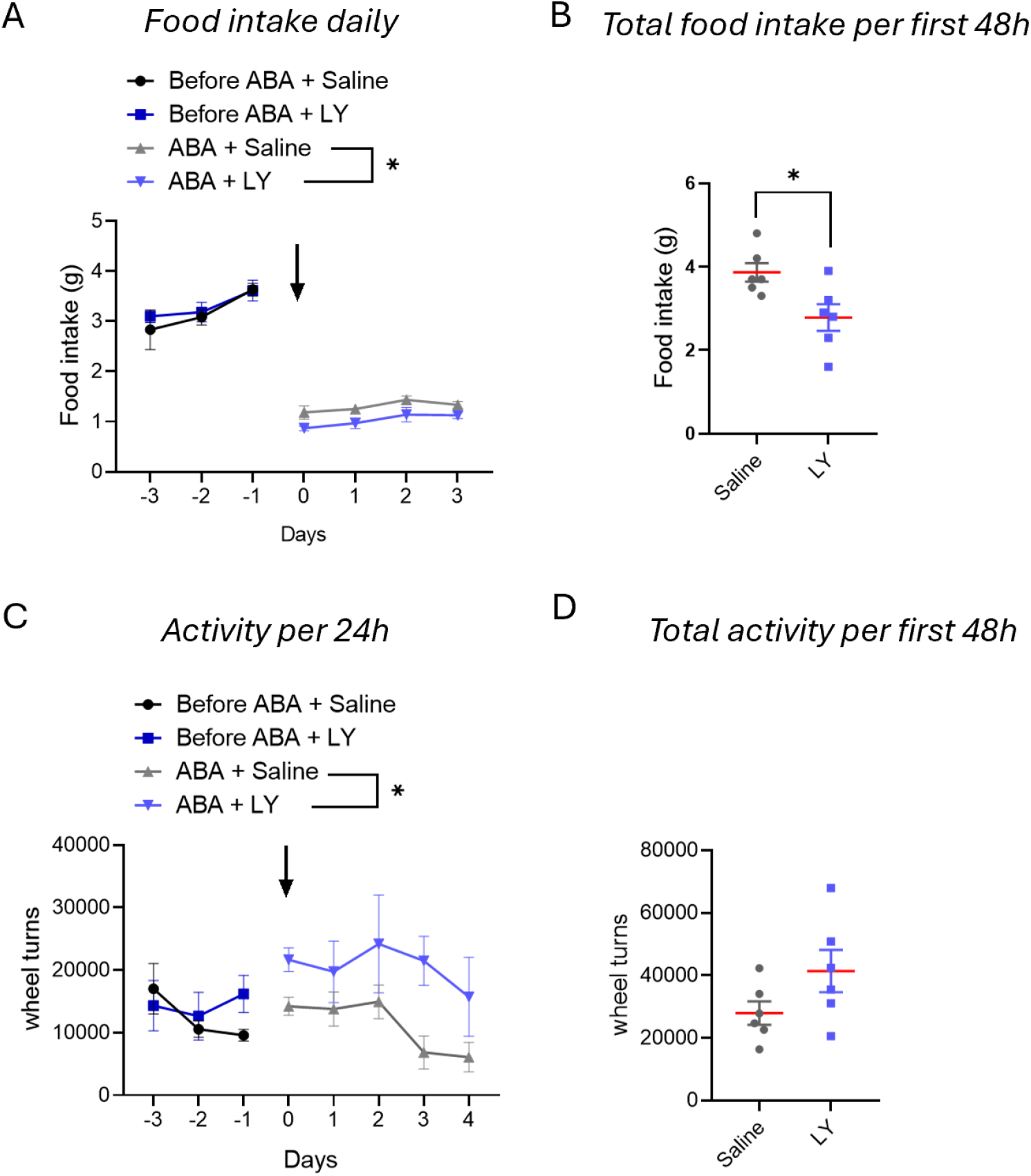
Effects of mGluR2/3 antagonism on feeding and activity behaviors in the ABA model. **A**. Food intake per 24 hours before the start of ABA and drug administration and per 1.5 hours at 24-hour intervals (days) for the “ABA” groups injected with saline or mGluR2/3 antagonist (LY). The start is marked by an arrow (6 animals per group, mixed effect model (REML), group effect before treatment: F (1, 10) = 0.4005, p=0.5410; ABA + LY vs. + saline treatment: F (1, 10) = 8.056, *p= 0.0176, data shown as the mean ± S.E.M.). **B**. Total food intake per 48 hours in the “ABA” groups injected with saline or LY (saline vs. LY, t test, t=2.778, df=10, *p=0.0195, data shown as the mean ± S.E.M.). **C**. Activity (wheel turns) per 24 hours before the start of ABA and drug administration and for the “ABA” groups injected with saline or mGluR2/3 antagonist (LY). The start is marked by an arrow (N=6, two-way ANOVA, group effect: F (1, 10) = 0.2798, p=0.6084; LY effect: F (1, 10) = 6.708, *p=0.027, data shown as the mean ± S.E.M.). **D**. Total activity (wheel turns) per 48 hours in the “ABA” groups injected with saline or LY (N=6, t test, t=1.736, df=10, p=0.1133, data shown as the mean ± S.E.M.).

## 4. Discussion

The results from our experiments point toward the manipulation of the metabotropic 2/3-glutamate receptors, be it by agonist or antagonist, to be detrimental to the animals in the ABA model at the acute stage of anorexia symptoms. The main effects of both pharmacological treatments on food intake and body weight for the ABA model emerged within 48 hours of daily administration. Although the mGluR2/3 agonist reduced food intake in mice with restricted feeding time, it did not affect their body weight as detrimentally as it did for the ABA model within 48 hours. There was no effect of the treatments on body weight in the mice fed ad libitum. Previously, metabotropic 2/3-receptor antagonism was shown to decrease food intake after overnight food deprivation in male rats (Semenova and Markou, 2007). Our data align with this finding. Despite our expectation that pharmacological targeting with the agonist would increase food intake, it had the opposite effect. Not only did this not improve the condition, but it worsened it. The paradoxical action of both agonists and antagonists of mGluR2/3 on behavior has been demonstrated previously in depression models, where both treatments led to similar antidepressant-like behavioral outcomes (Chaki et al., 2013). At the behavioral level, the same outcome can occur when both manipulations converge on a shared downstream bottleneck, such as the same brain area or neuronal ensembles, even if the upstream routes differ, and when opposite manipulations result in their direct or indirect activation (due to disinhibition). For example, c-fos expression analysis in rats treated with an mGluR2/3 agonist and antagonist showed partial overlap in activated brain areas such as the paraventricular and centromedial nucleus of the thalamus, central amygdala, lateral parabrachial nucleus, and bed nucleus of the stria terminalis (Hetzenauer et al., 2008; Linden et al., 2005).

Neurotransmitter dysfunction, particularly of glutamate, has been implicated in AN pathophysiology (Godlewska et al., 2017; Hermens et al., 2020; Keeler et al., 2024; Mitchell et al., 2023). Several studies have indicated reduced levels of glutamate in the cortex of AN patients (Castro-Fornieles et al., 2007; Godlewska et al., 2017). At the functional level, patients with AN have been demonstrated to show abnormal thalamocortical connectivity (Biezonski et al., 2016). Previous studies have shown hypermetabolism in the thalamus (Takano et al., 2001) as well as in the frontal lobe, hippocampus, amygdala, and insula (Zhang et al., 2013), where the major neuronal component is glutamatergic. Findings in AN patients have aligned with observations in animal models. Substantial neuronal activation and hypermetabolism were found in the thalamic nuclei of ABA rodents (Schéle, 2024; van Kuyck et al., 2007). Strong amygdala activation was also demonstrated (Mottarlini et al., 2022a). Glutamatergic synaptic malfunction was found in the hippocampus, NAc, and mPFC (Chen et al., 2017; Mottarlini et al., 2024, 2022b, 2020).

Ketamine demonstrated a beneficial effect on the ABA model in several studies (Dong et al., 2025b, 2025a; Dong and Aoki, 2025b; Li et al., 2024) and is considered to be a promising candidate for anorexia treatment (Hermens et al., 2020). Antidepressant-relevant actions of ketamine converge with mGluR2 signaling (Zanos et al., 2019). Negative modulation of mGluR2/3 rescues normal thalamocortical connectivity in the model of depression, suggesting overlap in the final common pathway between receptor antagonism and ketamine action (Joffe et al., 2020). On the other hand, the agonism of mGluR2/3 can promote NMDA receptor function (Rosenberg et al., 2016), which would be the opposite action compared to ketamine, which shows antagonism toward the NMDA receptor (Mitchell et al., 2023). Despite this overlap in the pharmacological functional pathways, neither the mGluR2/3 agonist nor the antagonist alleviated acute anorexia symptoms in the ABA model.

Overall, the data additionally indicate a contribution of glutamatergic transmission to anorexia symptoms. Although the mechanism by which mGluR2/3 can produce such a detrimental effect on the ABA phenotype is not known and was not described in our study, targeting mGluR2/3 may therefore be insufficiently specific in the context of anorexia-related circuit dysfunction, where both increased and decreased glutamatergic tone can be harmful. Anorexia nervosa continues to resist direct weight-affecting pharmacological treatments (Bauschka et al., 2023) and remains a complex psychiatric condition, with the main possible intervention focus on treating comorbidities and symptoms such as anxiety and obsessive compulsions. The II group of glutamate receptor agonists and antagonists could still have possible use for testing as treatments in preclinical models for other eating disorders, such as binge eating disorder (BED), or for obesity, with their actions distinct from those of GLP-1 analogs.

## 5. Conclusions

Pharmacological targeting of mGluR2/3 has a fast, detrimental effect on anorexia symptoms in the ABA model, affecting food intake, body weight loss, and survival. Despite possible convergence in action with ketamine, the mGluR2/3 antagonist does not improve the condition of animals with induced anorexia but worsens it. Surprisingly, the opposite action of the agonist produces similar behavioral and physiological effects as the antagonist. Further investigations are warranted to find a mechanism by which these effects are associated with glutamatergic transmission.

## Abbreviations

ABA: activity-based anorexia
AMPA: α-amino-3-hydroxy-5-methyl-4-isoxazolepropionic acid
GLP-1: Glucagon-Like Peptide-1
mGluR: metabotropic glutamate receptor
mGluR2/3: metabotropic glutamate receptor 2/3
mPFC: medial prefrontal cortex
NAc: nucleus accumbens
NMDA: N-methyl-D-aspartate
REML: Restricted Maximum Likelihood
RM ANOVA: Repeated measures Analysis Of Variance
SSRIs: Selective serotonin reuptake inhibitors.

## CRediT authorship contribution statement

Tuomas Varjelus: Writing – original draft, review & editing, Investigation, Conceptualization, Formal analysis, Data curation. Atte Oksanen: Investigation, Formal analysis, Data curation; Laura Kaljala: Investigation, Data curation. Maria Ryazantseva: Writing – original draft, review & editing, Formal analysis, Data curation, Conceptualization, Visualization, Supervision. Teemu Aitta-aho: Writing – review & editing, Conceptualization, Supervision, Resources, Project administration, Funding acquisition.

## Acknowledgment

The authors thank the personnel of the Laboratory Animal Center of the University of Helsinki.

